# Beyond guard cells: Epidermal anatomy determines transient stomatal opening following rapid changes in hydration status

**DOI:** 10.1101/2024.02.15.580521

**Authors:** Maryam Alsadat Zekri, Daniel Tholen, Lukas Koller, Klara Voggeneder, Ingeborg Lang, Guillaume Theroux-Rancourt

## Abstract

The rapid and transient increase of stomatal conductance in response to a sudden water deficit in plants, known as the wrong-way response (WWR), is hypothesized to result from rapid turgor loss of epidermal cells, which pulls the guard cells apart, leading to the opening of the stomatal pore. Variations in WWR amplitude and duration could stem from differences in epidermal anatomy and/or cell density, which influence the localized turgor responses driving stomatal movement. This study tested this hypothesis by aiming to induce WWRs through both a sudden increase in the leaf-to-atmosphere vapor-pressure deficit and leaf excision. We investigated the WWR kinetics in five *Arabidopsis thaliana* lines with varying epidermal characteristics. Stomatal conductance was measured at high temporal resolution using infrared gas analysis and epidermal anatomical traits were characterized using scanning electron microscopy. Interestingly, no WWR was detectable following a rapid decrease in air humidity. By contrast, leaf excision triggered a WWR across the majority of leaves examined, enabling robust analysis of response kinetics. While variation in magnitude and duration of the WWR among different lines was small, the duration and the amplitude of the WWR was strongly correlated with pavement cell size. No association between these WWR metrics and stomatal traits or shape-related characteristics such as cell eccentricity or lobe density were detected. These findings indicate that pavement cell size is a key determinant of the mechanical coupling underlying the WWR.

## Introduction

Stomata are tiny pores on the leaf surface that act as gatekeepers between the atmosphere and the leaf interior, regulating the intake of CO₂ and limiting transpiration. These pores are highly sensitive to environmental conditions that alter the plant’s water potential, such as soil drought or declining air humidity. Stomatal closure in response to a water deficit is typically explained by a hydroactive feedback mechanism involving the accumulation of abscisic acid (ABA), which triggers the release of osmotic solutes from the guard cells. This process reduces osmotic pressure in these cells, leading to the closure of the stomata (Buckley, 2019; Bharath et al., 2021). Nevertheless, stomatal closure is not the immediate response of plants to a rapid decline in water potential. Instead, a transient opening of the stomata immediately after the drop in water potential and before the eventual closure of stomata is typically observed (Darwin, 1898; Ivanov, 1928; Buckley et al., 2011; Cadart et al., 2019). This phenomenon, known as the wrong-way response (WWR), is considered a hydropassive feedback resulting from reduced epidermal turgor pressure, which pulls apart the turgid guard cells, causing stomata to open (Cowan, 1972; Buckley et al., 2011; Cadart et al., 2019). The role of pavement cells in pulling apart the guard cells following an immediate reduction in water potential has been described as the epidermal mechanical advantage (Buckley, 2019). This concept implies that the transient opening of a stoma in response to the drop in water potential is influenced not only by the physical properties of the stoma itself, but also by those of the surrounding pavement cells. While it is well-documented that physical properties of stomata, such as size and density, play a significant role in their movement (Franks & Beerling, 2009; Buckley et al., 2011; Carlson et al., 2015; Bertolino et al., 2019; Haworth et al., 2021), there remains a gap in our understanding of how physical characteristics of stomata and pavement cells affect the dynamics of transient opening. With our study we aim to address this gap by providing empirical data to shed light on the hypothesis of the epidermal mechanical advantage.

The role of pavement cell geometry in stomatal development (Dow and Bergmann, 2014) and responses to environmental stresses (Clauw et al., 2015) suggests that epidermal anatomy is a significant factor that modulates responses of plants to variations in water potential. In addition, smaller cells with a higher surface-area-to-volume ratio respond faster to osmotic changes (Drake et al., 2013; Giday et al., 2013; Kardiman and Rabild, 2017; Nunes et al., 2020) and it has been suggested that -mechanical pressures impact smaller structures less significantly than larger ones (Bassel et al., 2014). In this study, we investigated the kinetics of the WWR in five *Arabidopsis* lines with contrasting epidermal anatomical features. Pavement cell characteristics such as area, eccentricity, and the size and number of lobes were analyzed to elucidate their role in the hydropassive response. We hypothesized that *Arabidopsis* lines with smaller pavement cells and stomata would be more resilient to mechanical forces resulting from water potential perturbations. The observed differences in pavement cell characteristics highlight the role of epidermal anatomy in leaf responses to water stress, suggesting that selecting or modifying these traits could improve plant water-use efficiency and stress tolerance.

## Material and Methods

### Plant material and growth conditions

Seeds of five *A. thaliana* lines were chosen based on their variation in leaf epidermal traits and provided by the Nottingham Arabidopsis Stock Centre (NASC): Col-8 (NASC accession: N60000) of Columbia background, was chosen as the wild-type control; in the *epf1epf2* double mutant (NASC N67139), the loss of EPF1 and EPF2 drives excessive entry of epidermal cells into the stomatal lineage, resulting in a lower average epidermal cell area and a high proportion of small, unspecialized epidermal cells across the tissue, in addition to exhibiting elevated stomatal density (Hara et al. 2007, 2009; Hunt and Gray, 2009); *lcd1-1* (N6359) was reported to have smaller leaves and a higher fraction of epidermal tissue (Gonzalez-Bayon et al. 2006), but may also have a lower cell density in the mesophyll (Barth and Conklin 2003); UBP (UBP15-overexpression: N70771) has been shown to have increased epidermal cell numbers (Liu et al. 2008); and SALK069 (SALK_069127: N569127) was reported to have larger epidermal cells (Jalakas et al. 2018).

14 plants per line were grown in individual pots (diameter = 13 cm) containing 50% potting soil (Kranzinger GmbH, Austria), 33% quartz sand, and 17% perlite. Plants were grown for five weeks in a growth room (Walk-in FytoScope, Conviron MTPS 120, Manitoba, Canada) with a 16 h photoperiod, 130-150 µmol m^-2^ s^-1^ photosynthetic photon flux density, 22/18°C day/night temperature, and 60-65% humidity (RH). Only well-grown and fully expanded leaves were used for further analysis.

### Stomatal density and pore length

To extract features of epidermal cells, detailed images of the epidermis were collected by Scanning Electron Microscopy (SEM). Talbot and White (2013) compared several different tissue preparation methods for SEM and found that fixation by methanol and dehydration in ethanol typically results in the best preservation of tissue size and morphology (Talbot and White, 2013). We therefore adapted this method for our purposes. 10 mature leaves per line were fixed in 100% methanol for 10 minutes, followed by two dehydration steps in 100% ethanol for 30 minutes. Critical point drying (Leica, CPD300) was performed based on the manufacturer’s protocol (CPD300, 2012). Three leaves were randomly selected from each line to collect anatomical features. Two pieces were excised from each leaf, one representing the abaxial surface and the other the adaxial surface and mounted on stubs using double-sided adhesive carbon tape. The leaf pieces were cut from the well-expanded portion of the leaf, avoiding the edges, tip, areas near the main vein, and petiole. Since the imaging was performed under high vacuum, the samples were coated with gold (Small Multi-target Metal Ion Sputtering Coater Machine, Model Number, TMAX_BY_JS11). Three images at different locations on the abaxial and adaxial surface were acquired by SEM (JEOL IT300) (Fig. 1). On each image, three individual squares (5000 x 5000 µm) were selected to measure stomatal density. Stomatal length was defined as the length of the guard cells (Savvides et al., 2011) and was measured on 100 stomatal apertures per leaf using ImageJ (Schindelin et al., 2012).

**Figure 1.**
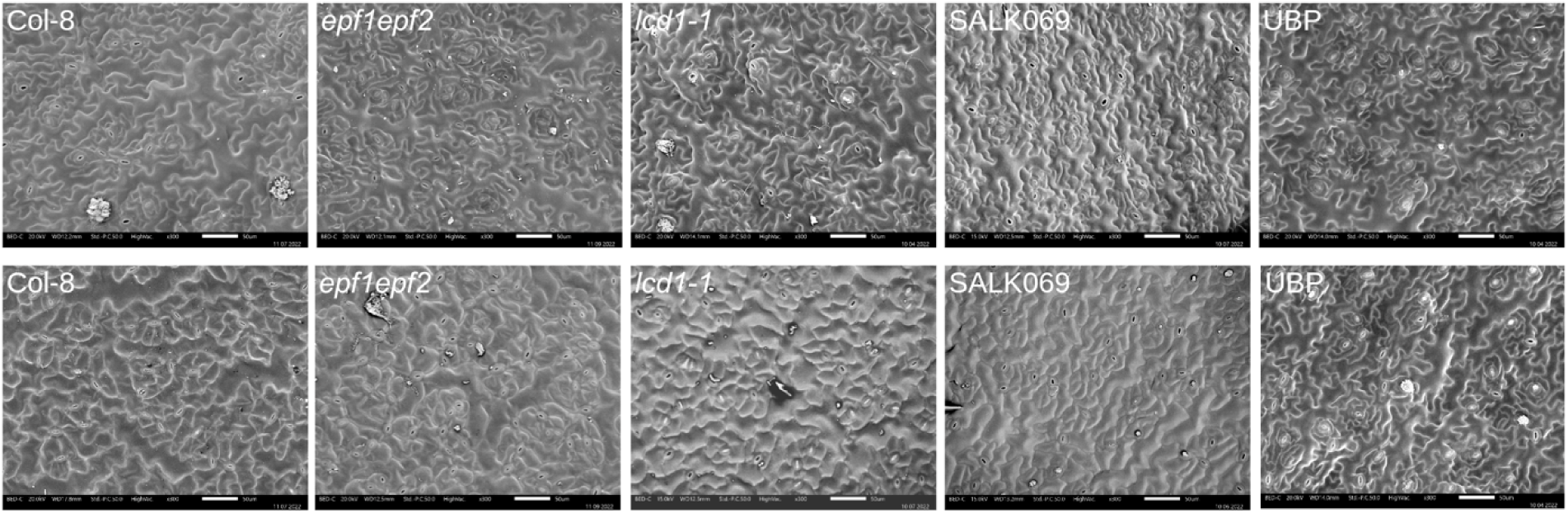
SEM images of abaxial (upper row) and adaxial (lower row) surfaces of five *Arabidopsis* lines. Scale bar: 50µm.

### Extraction of epidermal cell features

Pavement cell morphology was quantified using the PaCeQuant software platform (Möller et al., 2017), and our analysis expanded upon the structural trait framework described by Sapala et al. (2019). Because cell size was our primary focus, our evaluation centered on size-related metrics such as cell area and the longest path length (the longest distance between two points measured entirely within the cell’s boundary). We also included several morphological traits that were expected to covary with pavement cell growth: total lobe count per cell, average lobe length (size of protrusion measured on the cell boundary), branch length (lobe length measured as internal skeleton segment length in PaCeQuant), cell circularity, and convexity. To capture geometry independent of cell size, we measured eccentricity (quantifies cell elongation), lobe density (lobes per convex hull perimeter), and average lobe width at the midpoint.

To extract features of pavement cells, SEM images were analysed using the PaCeQuant plugin of Fiji/ImageJ (Möller et al., 2017). Since this plugin requires high-contrast images, cell boundaries on SEM images were first manually highlighted in GIMP-2.10 (The GIMP Development Team, 2019) and added to the image as a second layer. This layer was then extracted and used for the analysis of the pavement cells using the PaCeQuant plugin of Fiji/ImageJ (Möller et al., 2017; see Supplemental Figure S1 for workflow). Figure 1 shows representative SEM images of the abaxial and adaxial epidermis of the five *Arabidopsis* lines.

### Gas exchange measurements and WWR experiments

The plants were watered daily for a cultivation period of six weeks. After the growth period, stomatal conductance to water vapor (*g_sw_*) was measured using a portable infrared gas analyzer system (LI-6800, LI-COR Biosciences, Lincoln, NE, USA). The leaf temperature inside the chamber was maintained at 25°C along with a light level of 400 μmol m^−2^ s^−1^ and a reference CO_2_ concentration of 400 µmol mol^-1^. Air flow rates through the sample chamber were kept constant at 400 µmol s^-1^ during leaf excision but increased to 500 µmol s^-1^ during humidity changes to facilitate faster mixing of air inside the chamber. The humidity of the air entering the leaf chamber was set to a relative humidity of 60%, corresponding to a leaf-to-atmosphere vapor-pressure deficit (VPD) of 1.2 kPa. Leaf temperature was controlled by the LI-6800 and set to 25 °C throughout the experiments. Before initiating the WWR experiments, leaves were acclimated for 15 to 60 minutes under these conditions until steady-state gas exchange was reached, i.e. when fluctuations in *g_sw_* did not exceed 0.1 mol m^-2^ s^-1^ for 5 minutes. Subsequently, the humidity was reduced to ∼19% RH, corresponding to a VPD of 2.5 kPa. To study epidermal anatomy, 10 leaves per line were labeled and put aside for subsequent microscopic analysis (see below). Seven leaves were used for the second WWR experiment, in which their petiole was cut just below the lamina to separate the leaf from the plant using a razor blade (called “excision” hereafter).

### Correction for lags resulting from chamber mixing and water desorption

After changing leaf chamber humidity, the water differential fluctuates temporarily while the incoming air mixes throughout the chamber volume. Zait et al. (2024) characterized the response of artificial plastic leaves and concluded that water concentrations were unstable for 3 minutes post-switch due to the gas turnover time of the chamber. Because the effect of water desorption from chamber walls can persist even longer and bias transpiration measurements, Buckley et al. (2011) fitted a two-phase exponential model to the empty chamber response to correct for both mixing and sorption. We followed a similar approach by using a 200 µm Polytetrafluoroethylene foil (PTFE; LI-COR Biosciences, Lincoln, NE, USA) representing a leaf as control measurement. We subsequently fitted a smoothed biphasic power law to the relation between time and the difference between sample and reference humidity. Sample water concentrations were corrected using this model, and stomatal conductance was subsequently recalculated using the GasanalyzeR package (Tholen, 2024). Even with this correction, large outliers and fluctuations in leaf temperature were observed immediately after changing the VPD. Because the WWR peak is expected to occur only after several minutes (Buckley et al. 2011), we excluded data from the first 36 seconds post-switch to avoid such artefactual gradients.

### WWR kinetics

The kinetics of the WWR following a step increase in VPD or petiole excision were evaluated according to Buckley et al. (2011) using three kinetic parameters: amplitude (*S*), duration (*L*), and response rate (*V = S / L*). Duration (*L*) is the elapsed time between excision or VPD change and the moment stomatal conductance (*g*_sw_) begins to decline from its maximum value, corresponding to the time-to peak if a WWR is present. Amplitude (*S*) represents the relative increase in stomatal conductance (*g*_sw_) after duration *L*, calculated as (*g*_L_ - *g*_baseline_ ) / *g*_baseline_. When a peak was absent and *g*_sw_ declined immediately, *S* was clamped to zero.

### Statistical analysis

Differences between groups were tested using a one-way anova or linear mixed-effect model using R version 4.3.2 (R Core Team, 2024) and the lme4 package (Bates, 2015). Pairwise comparisons between means were done using the emmeans package (Lenth and Piaskowski, 2026) using the Tukey method for *p*-value adjustment. The normality of the data was confirmed using a Shapiro–Wilk test. We hypothesized that WWR metrics scale with pavement-cell size. Since different leaves were sampled, correlations were analyzed using genotype means using cell area as the main metric for size measure and eccentricity as a size-unrelated form measure, alongside stomatal density and size. Monotone associations were quantified by Spearman’s ρ with an exact permutation test over all 120 genotype relabellings (minimum attainable two-sided *p* for n=5: 0.017). To test whether cell area was associated with WWR metrics, the results were Holm-corrected across the two co-primary response parameters *S* and *L* (V is shown for illustration only as it is fully dependent on both other parameters).

## Results

### Variation in stomatal density, guard cell length, and pavement cell features

The mean stomatal density of *epf1epf2* was typically doubled relative to that of the other lines (p < 0.001; Fig. 2), and this effect was consistent on both adaxial and abaxial sides. There were no significant differences in total stomatal density among Col-8, *lcd1-1*, SALK069, and UBP.

**Figure 2.**
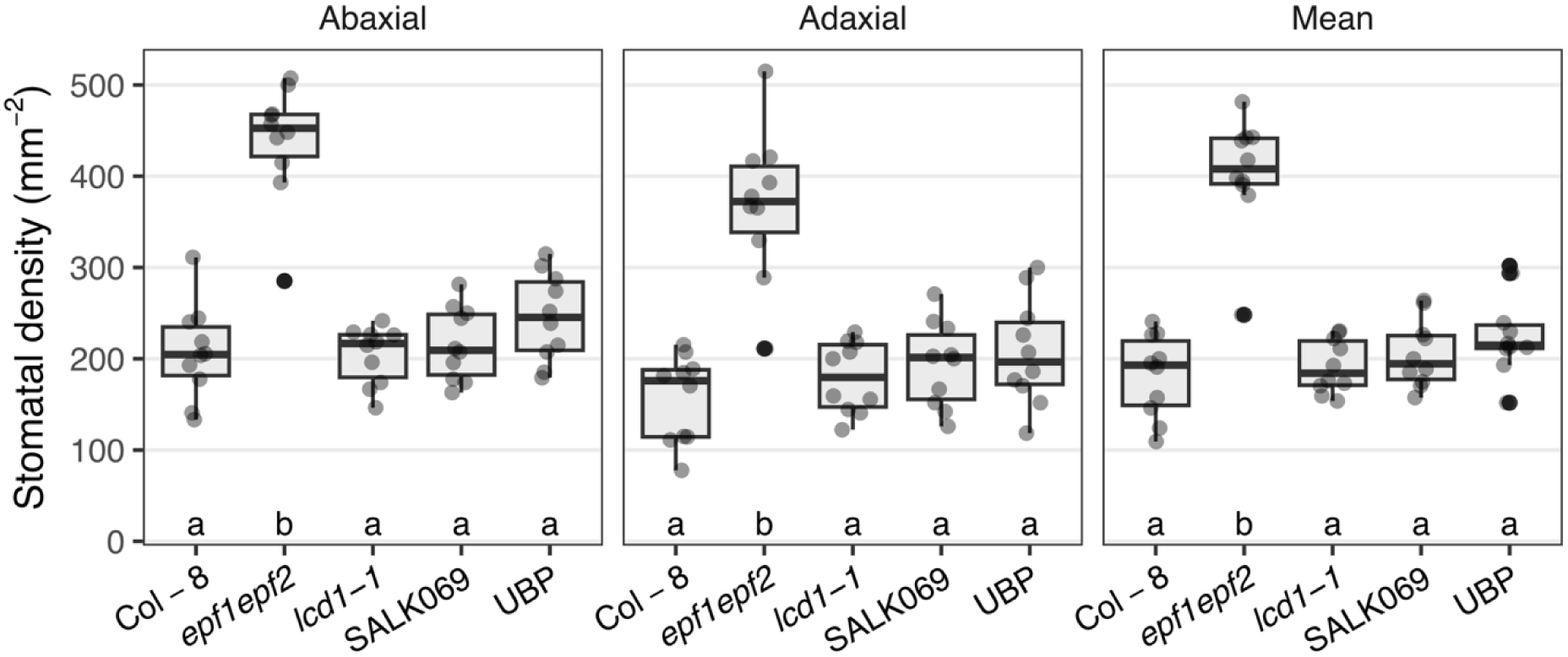
Stomatal density of 5 different *Arabidopsis* lines on the abaxial and adaxial surface, and in total. Different letters indicate a significant difference according to Tukey’s honest significant difference test (n = 10).

Guard cell length was generally consistent across lines, though UBP guard cells on both leaf surfaces were 5–7% longer than those of *lcd1-1* (*p* < 0.05). A similar increase was observed relative to *epf1epf2*, but this difference reached statistical significance only when data from the adaxial and abaxial surfaces were pooled (Fig. 3).

**Figure 3.**
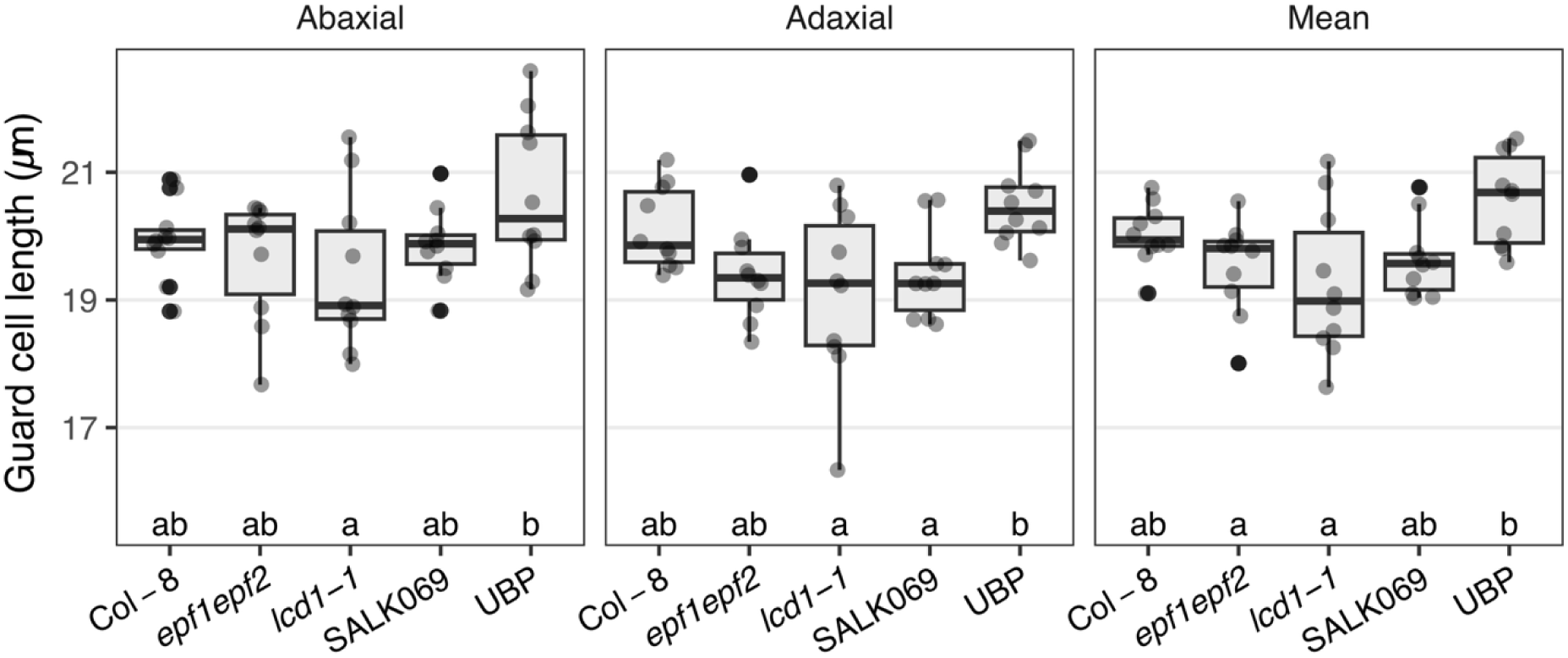
Size of abaxial and adaxial guard cells, and mean size of guard cells from both surfaces, among the five different *Arabidopsis* lines studied. Different letters indicate a significant difference according to Tukey’s honest significant difference test (n = 10)

Projected areas of the pavement cells on the abaxial side were largest for Col-8 and UBP, although the difference only achieved statistical significance in comparison with epf*1epf2* (*p* = 0.035 to < 0.0001; Fig. 4). *epf1epf2* cells on this leaf surface had shorter branch lengths compared to UBP (36%, *p* = 0.006) and marginally so when compared to Col-8 (27%, *p* = 0.075). The longest path through a cell was also shorter in *epf1epf2* than in Col-8 (49% shorter, *p* < 0.0001), SALK069 (40% shorter, *p* < 0.001), UBP (47% shorter, *p* < 0.0001), and *lcd1-1* (39% shorter, *p* < 0.001). Eccentricity was similar between all lines, indicating that pavement cells of all lines had similar elongation. *epf1epf2* and SALK069 had significantly thinner lobes than UBP (22% [*p* = 0.005] and 19% [*p* = 0.016] less, respectively), and *epf1epf2* also showed the lowest density of lobes, though only significantly different to *lcd1-1* (18% more, *p* = 0.017). *epf1epf2* pavement cells also showed higher circularity than Col-8 (32%, *p* < 0.001), *lcd1-1* (29%, *p* < 0.001), SALK069 (25%, *p* = 0.003), and UBP (23% higher, *p* = 0.006). *epf1epf2* had the highest cell convexity, being 7-9% (*p* = 0.001 to < 0.0001) higher than all other lines, and had 44-52% fewer lobes compared to the other lines (p < 0.0001). Results for the adaxial leaf side were very similar, although the differences were typically somewhat smaller (Supplemental Fig. S2).

**Figure 4.**
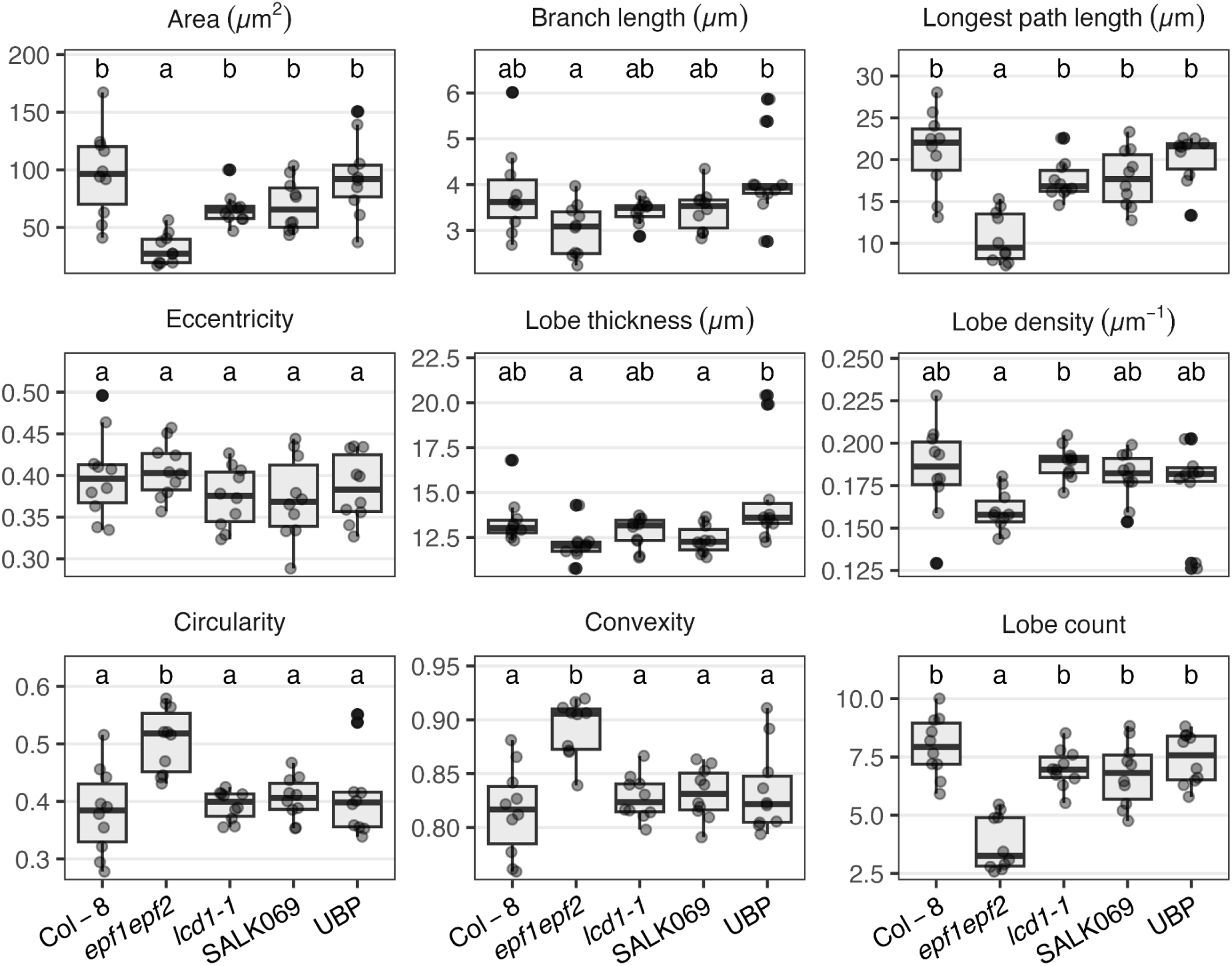
Anatomical characteristics of epidermal cells on the abaxial leaf surface in the five lines of *Arabidopsis*, presenting a selection of size-related traits (top row), size-independent traits (middle row), and additional geometrical traits that covary with size (bottom row). Different letters represent a statistically significant difference between the lines for each trait analyzed (n = 10).

### Genotypic variation in the WWR of Arabidopsis is pronounced after excision, but absent after a VPD shift

Gas-exchange data showed clear changes in water mole fractions and transpiration rates after a change in VPD and after leaf excision. However, after accounting for chamber mixing and water desorption, the increase in *g_sw_* in response to a VPD change was negligible in all five lines (Figure 5), as well as with using a PTFE foil for control (Supplemental Figure S3). Leaf excision induced a WWR in most leaves (Figures 5 & 6), though the magnitude of the peak is partially obscured in the genotype average due to slight variation in response kinetics among leaves. Aligning the data by normalizing both *g*_sw_ and time resolves the dynamics more clearly (Figure S4). The WWR occurred irrespective of previous VPD conditions or humidity transitions (Supplemental Figures S4 and S5).

**Figure 5.**
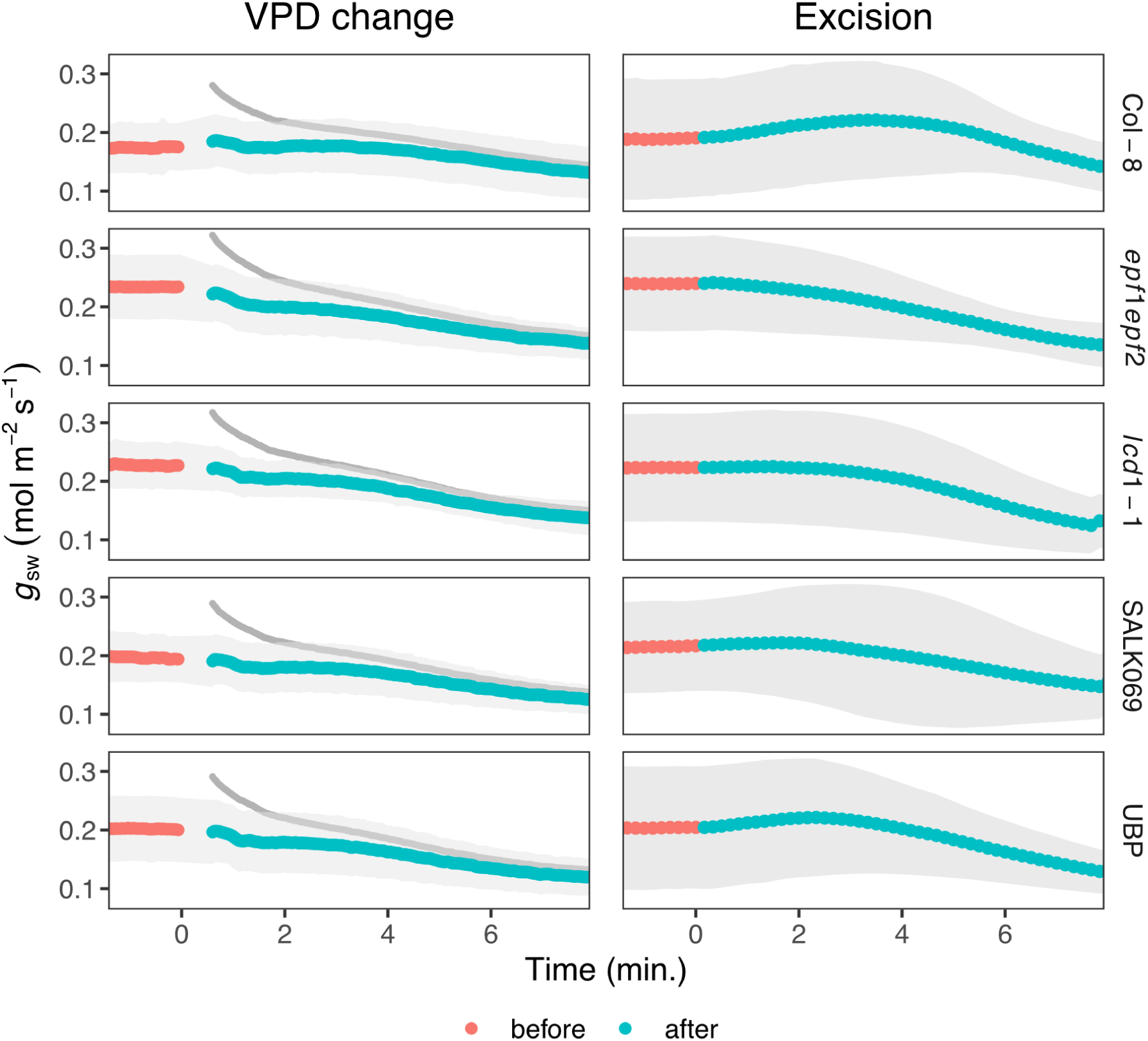
Stomatal conductance to water vapor (*g*_sw_) over time in five different Arabidopsis lines before and after changing the leaf to chamber air vapor-pressure deficit (VPD), and before and after cutting the petiole (excision). The average of 5-7 (excision) or 10-14 (VPD change) individual leaves was shown. The ribbon represents the standard deviation. Red points indicate measurements before the humidity switch (at a VPD of 1.2 kPa) or before excision (at a VPD of 2.5 kPa). Cyan points indicate measurements at a VPD of 2.5 kPa after changing chamber humidity or after excision. The grey lines in the VPD change column indicate the raw measurements without any correction for mixing and water desorption (see material and methods).

To characterize the WWR following leaf excision in the different lines, we quantified the relative amplitude (*S*), duration (*L*), and the rate of stomatal opening (*V* = *S*/*L*). Response characteristics were consistent across all 5 lines, with the exception of *epf1epf2, which* exhibited significantly lower *L* and *S* (Figure 6). Interestingly, several replicates did not show a WWR, increasing the variance and reducing the amplitude and duration of the averaged response (Figure 5). However, a Fisher’s exact test revealed no significant differences in WWR incidence among lines (p > 0.4 after correction). Substantial variability of the WWR response, including complete absence of the response on some leaves of individual plants, has been reported before (e.g. Buckley et al. 2011, Manandhar et al. 2026).

**Figure 6.**
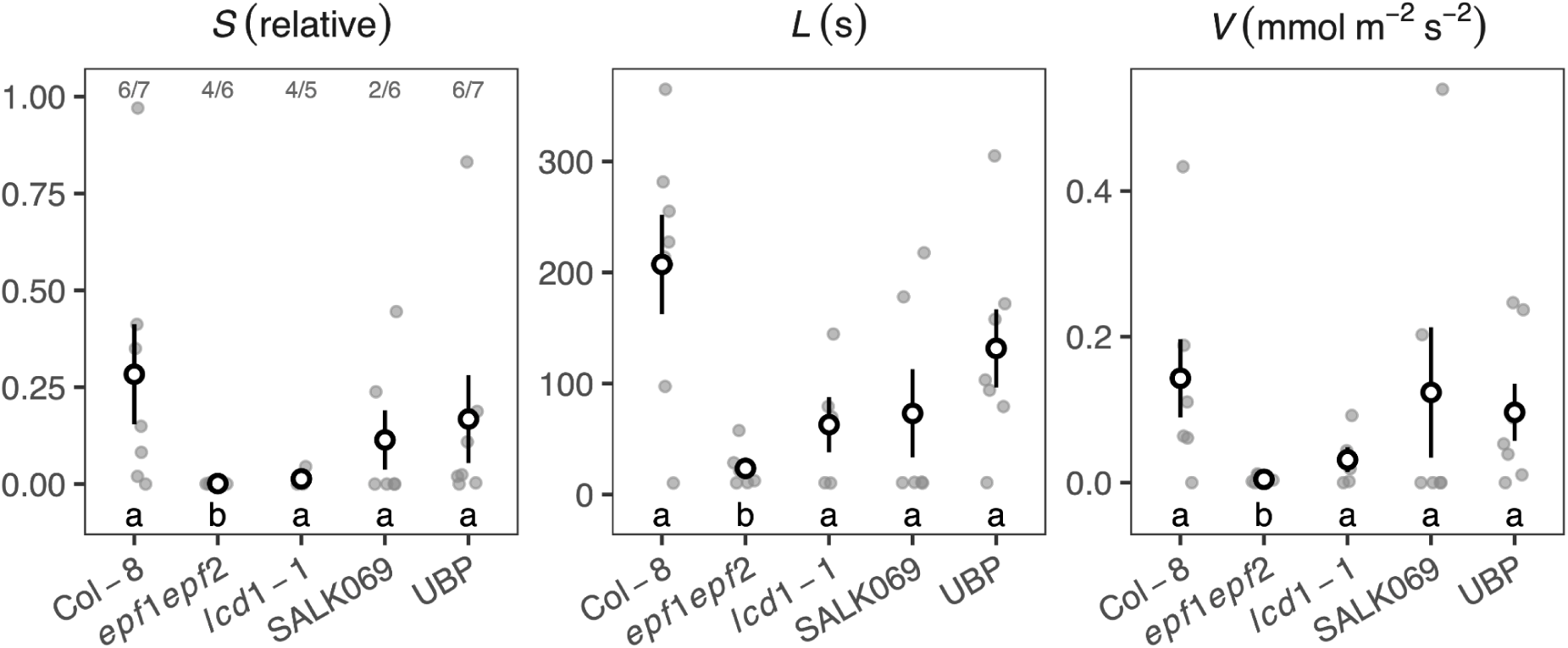
WWR parameters in response to leaf excision (n= 5-7). *S*) amplitude of the WWR, relative to steady-state stomatal conductance before excision. *L*) duration of the WWR. *V*) rate of the WWR, computed as *S*/*L*. Different letters represent a statistically significant difference between the lines for each trait analyzed. The ratios above each line in panel *S* indicate the number of responsive leaves out of the total number of leaves tested.

### Pavement cell size traits correlate with WWR amplitude and duration

WWR amplitude (*S*) and duration (*L*) were strongly monotonically associated with cell size, whether measured by area or longest path length (Spearman ρ = 1 for both; Holm-adjusted *p* = 0.02). This relationship remained invariant when combining size metrics into an eleven-trait composite index (Supplemental Figure S6). In contrast, two shape descriptors that were geometrically independent of size, eccentricity and lobe density, showed no association with the physiological traits (ρ ≤ 0.4). Furthermore, an exploratory analysis of all PaCeQuant metrics revealed no further associations with other shape-related traits (Supplemental Figure S6). Similarly, no significant association was observed between the physiological responses and stomatal density (ρ = -0.6, *p* = 0.35) or guard cell length (ρ = 0.8, *p* = 0.13; Figure 7).

**Figure 7.**
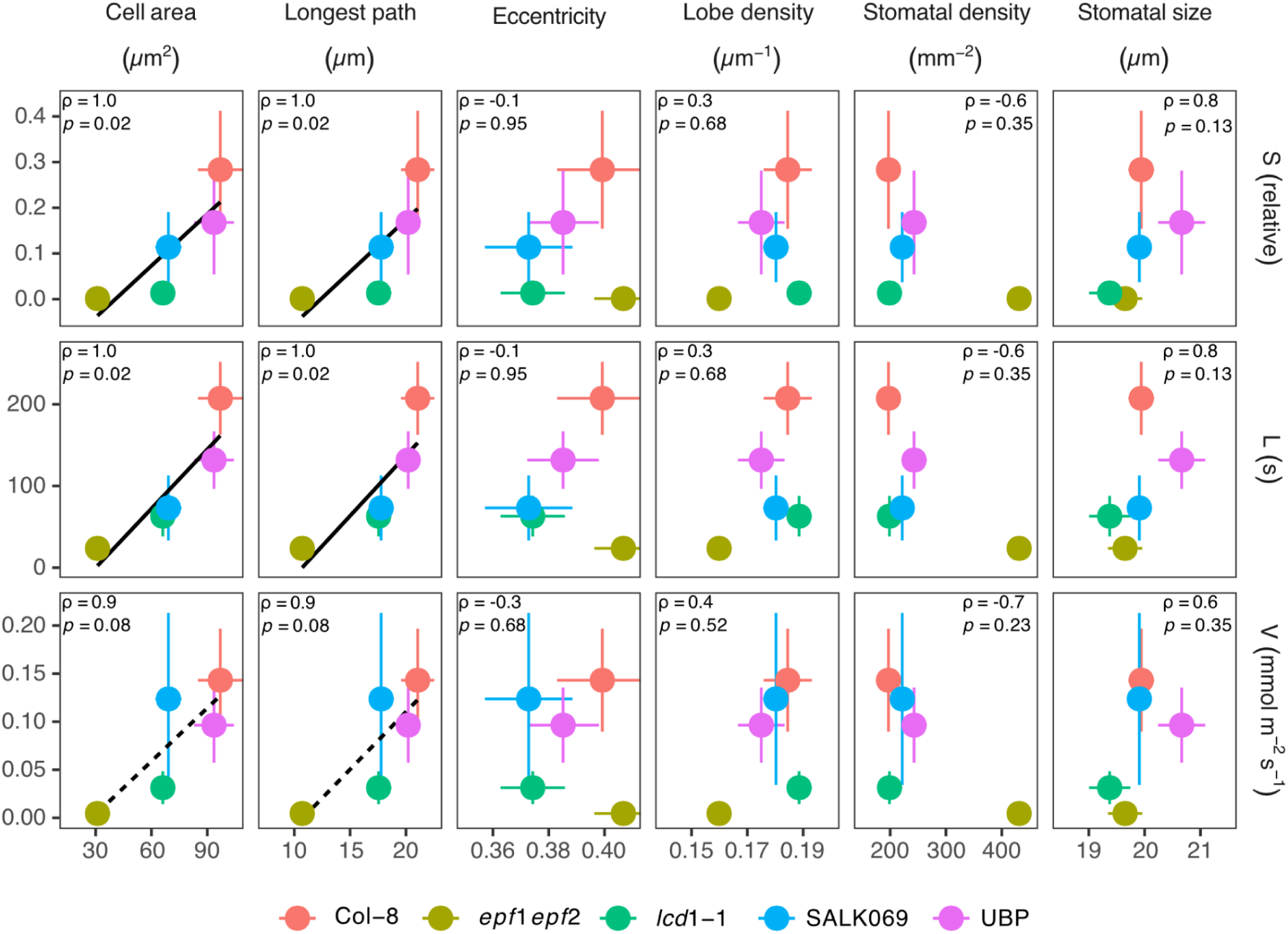
Relationships between the relative amplitude (*S*, top row), the duration (*L*, middle row), and the rate (*V*, bottom row) of stomatal transient opening following leaf excision and anatomical properties on the abaxial surface for five studied *A. thaliana* lines. Points show mean values for each *Arabidopsis* line, with horizontal and vertical bars representing standard deviation for each variable. Panels display Spearman’s ρ and *p*-values derived from exact permutation tests. Continuous lines *p* < 0.05, dashed lines *p* < 0.1.

## Discussion

In this study, we analyzed sudden stomatal opening in response to a drop in water potential triggered either by leaf excision or by a switch from low to high VPD, commonly referred to as the wrong-way response (WWR). The most widely accepted hypothesis explaining the WWR suggests a passive and ABA-independent pathway following a decrease in water potential: a decline in turgor pressure of the pavement cells pulls the guard cells apart, resulting in transient stomatal opening (Buckley, 2019).

Buckley et al. (2011) observed a WWR after increasing VPD that peaked between 7 and 11 minutes after changing humidity in 20 species with heterobaric or homobaric leaves and herbaceous or woody life forms, though they did not consistently observe transient responses to humidity in all individuals or species. For different *Arabidopsis* lines, Zait et al. (2024) reported a WWR following a humidity switch that peaked within 3 min, with the total period of elevated *g*_sw_ lasting up to 17 min. At our flow speeds (500 µmol s^-1^), the effect of gas-mixing and water desorption from the chamber walls can persist for 10 min after a change in chamber humidity. Since water desorption can artefactually increase transpiration, it was corrected for using control measurements adapting the methodology described by Buckley et al. (2011). Following this correction, we found no evidence for an increase in *g_sw_* after a humidity shift in any of the *Arabidopsis* lines studied here (Figure 5). Under CO_2_ limitation, transient stomatal opening should elevate net CO_2_ assimilation alongside *g*_sw_. Thus, measuring CO_2_ dynamics offers a robust, independent verification of the WWR because the CO_2_ flux is free from the chamber wall sorption effects that complicate water vapor measurements. Accordingly, we observed a distinct post-excision peak in assimilation rate, whereas VPD shifts elicited no detectable peak in assimilation rate (Supplemental Figure S7).

Our results highlight that failing to account for the effects of mixing and water desorption can easily lead to misinterpretation of the amplitude and duration of the WWR, and that these corrections are necessary when imposing rapid changes in VPD: otherwise, results should be viewed as relative responses, as some authors have noted (Hsu et al., 2021). We confirmed our results in separate experiments with the same *Arabidopsis* lines grown in different growth-chambers (with a photoperiod of 10 h and a light intensity of 300 µmol m^-2^ s^-1^) and measured with different gas-exchange equipment (GFS-3000, Heinz Walz GmbH, Effeltrich, Germany). Since it takes time for the humidity concentration in the gas-exchange chamber to stabilize, a response faster than 36 seconds could have occurred. However, it is very unlikely that the hydroactive response would trigger after such a short timeframe and suppress a WWR (Buckley et al., 2011; Vialet-Chabrand et al., 2017).

Increasing VPD instantly accelerates evaporation, which rapidly cools the leaf via latent heat loss. A thermocouple registers this cooling within seconds, causing the LI6800 software to calculate a lower internal saturation vapor concentration (*w*_i_) and a smaller vapor gradient (*w*_i_-*w*_a_). In contrast, the measured transpiration rate (*E*) relies on flushing air through the cuvette, creating an unavoidable time lag due to chamber mixing volume and flow rate. Because *g*_sw_ ≅ *E* / (*w*_i_-*w*_a_), this generates an artificial, temporary spike in the estimated conductance that closely mimics a transient wrong-way opening response. Keeping leaf temperature constant, as we did, limits this rapid change in VPD-induced evaporative cooling in amplitude (< -0.5°C) and in time (30-60 s; see Supplemental Figure S8). This period of time should also be seen as unreliable and matches the duration of the water sorption effect. We recommend that researchers investigating humidity-induced WWR tightly control and explicitly report leaf temperature dynamics. The absence of a VPD-induced WWR after correction for water sorption and under constant leaf temperature indicate that, under our growth and measurement conditions, the epidermal cells of *Arabidopsis* do not dehydrate rapidly enough following a humidity drop to trigger a WWR.

Leaf temperature remained unchanged following excision (Supplemental Figure S8) and we detected clear differences in the WWR among the tested lines, which were present under both high and low humidity levels before and after the VPD shift (Supplemental Figure S5). The line with the smallest pavement cells, *epf1epf2*, exhibited the shortest WWR (Figure 6). *S* and *L* were strongly correlated to the size of the pavement cells on the abaxial surface (through cell area and the longest path length within cells; Figure 7). Although PaceQuant yields numerous geometrical shape descriptors, none exhibited an unadjusted correlation with WWR metrics (Supplemental Figure S6). Although with five genotypes the statistical power for trait screening is limited, the clear separation between size and shape related metrics suggest that it is the size, rather than cell geometry, that dictates WWR dynamics. Thus, smaller cells may exert less backpressure on the guard cells, leading to a suppressed WWR. In agreement with this view, Bassel et al. (2014) demonstrated that when large embryonic cells of *Arabidopsis* sp. are subjected to pressure, they expand more than smaller cells. Similarly, results by Sapala et al. (2018) and Bidhendi et al. (2019) supported the view that plants confront mechanical stresses through higher cell interdigitation or smaller cell size. This implies that small cells are more resilient to volume changes under mechanical stresses and supports the notion that small structures are better able to confront mechanical stresses (Drake et al., 2013; Kardiman and Rabild, 2017; Théroux-Rancourt et al., 2021; Caine et al., 2023).

The observed differences in stomatal responses following the humidity or excision treatments may be related to the nature of the induced water deficit. The decrease in leaf water potential following low humidity is likely more gradual and primarily driven by external environmental factors. The water loss of epidermal cells is influenced by mechanical barriers such as the cuticle (Riederer and Schreiber, 2001), trichomes (Zhu et al., 2024), and, more generally, by the boundary layer conductance (Schuepp, 1993; Jones, 1999). Leaf excision, on the other hand, disrupts the water supply from the soil, causing a severe and abrupt decline in leaf water potential. Although epidermal properties may still affect the rate of dehydration, the latter may also be affected by other tissues in the hydraulic pathway, such as the bundle sheaths cells, which are known to control water flow from the xylem into the mesophyll cells (Buckley et al., 2011; Sade et al., 2015; Hua et al., 2021). In addition, embolism formation in the xylem may also influence the dynamics of the WWR (Salleo et al., 2000; Buckley et al., 2011). Therefore, the absence of a WWR following a change in humidity may be related to the capacity of the leaf to maintain the hydration status of the epidermis through continued water supply from the mesophyll or xylem. Buckley et al (2011) suggested that a low mesophyll elastic modulus and low hydraulic resistance between the bundle-sheath and epidermis dampen the WWR. *Arabidopsis* leaves likely exhibit such hydraulic traits. While this mechanism may prevent a WWR in response to a humidity shift, it may be insufficient to prevent the response to a more severe water deficit following leaf excision.

Our study focused on pavement cell geometry and with the exception of the *epf1epf2* mutant, stomatal size and density not differ significantly among our lines, distinguishing our dataset from studies explicitly targeting stomatal characteristics (Dow et al., 2014; Franks et al., 2015; Monda et al., 2016). Although shorter WWR durations and amplitudes were associated with smaller pavement cells, comparisons across distinct genotypes can be influenced by line-specific background traits. For example, SALK069 has impaired ABA catabolism (Okamoto et al. 2009). However, the WWR is widely accepted as an ABA-independent hydropassive response (Buckley 2019) and previous work in poplar showed that neither exogenous nor endogenous ABA affects WWR following leaf excision (Théroux-Rancourt, 2014). Additionally, while altered mesophyll cell density in *lcd1-1* (Barth & Conklin, 2003) may affect its steady-state gas exchange, it did not seem to deviate the correlations with epidermal cell size observed in our study (Figure 7), suggesting that WWR kinetics are driven more by localized guard cell turgor loss and/or epidermal hydraulics than by mesophyll structure.

The findings of this study enhance our understanding of stomatal responses and the mechanisms by which plants react to changes in water potential. Our results highlight the distinct roles of pavement cell geometry and stomatal anatomy in epidermal organization and demonstrate how cell packing specifically shapes leaf physiological responses. Understanding these relationships is essential for breeding programs aimed at improving plant resilience to environmental stressors. In addition, our results highlight how smaller epidermal cells allow leaves to reach steady-state gas exchange more quickly under fluctuating environmental conditions, suggesting tighter hydraulic coupling (Buckley et al. 2011). As herbaceous plants generally have lower hydraulic resistance than woody plants (Buckley et al. 2011), smaller epidermal cell size could further enhance hydraulic safety. Hence, both epidermal cell properties and within-leaf hydraulic compartmentalization (e.g. Buckley et al. 2011, Zwieniecki et al. 2007) govern responses to rapid environmental changes. Future studies should address how these transient events are regulated through the interplay between epidermal, mesophyll, and hydraulic leaf architecture.

## Supporting information

Workflow used to segment pavements cells. Using a SEM image (A), pavement cell delimitations were drawn using a graphic tablet (B). These hand-labeled

Anatomical characteristics of epidermal cells on the adaxial leaf surface in the five lines of A. thaliana. Different letters represent a statisticall

Stomatal conductance to water vapor (gsw) over time following a change in VPD from 1.2 kPa to 2.5 kPa for an empty gas-exchange chamber, for a chamber

Stomatal conductance to water vapor (gsw) relative to the value at the moment of excision in five Arabidopsis lines following leaf excision. To adjust

## Authors contribution

GTR, DT, IL, and MAZ conceived the project. MAZ, GTR and KV acquired gas exchange data. MAZ and LK acquired imaging data. MAZ, DT, and GTR analyzed the data. MAZ, DT, and GTR wrote the manuscript, with contributions IL. All authors agreed on the final version.

## AI generative statement

No AI tool was used to generate scientific content, analysis, interpretations, or results. All text was revised and edited by the authors, who take full responsibility for the accuracy and integrity of the content. The authors used the generative AI tool ChatGPT (OpenAI) for grammar and language improvement. The authors confirm that there is no AI authorship or contribution to the scientific conclusions of the paper.

## Data availability statement

The data that support the findings of this study are available on request from the corresponding authors.

## Acknowledgement

Microscopy (SEM Image acquisition) was performed at the Core Facility Cell Imaging and Ultrastructure Research, University of Vienna - member of the Vienna Life-Science Instruments (VLSI) and we are grateful to Daniela Gruber for guiding in SEM image acquisition. We gratefully acknowledge the funding from the Vienna Science and Technology Fund (WWTF) [10.47379/LS19013].

## Supplementary Information

**Figure S1.**
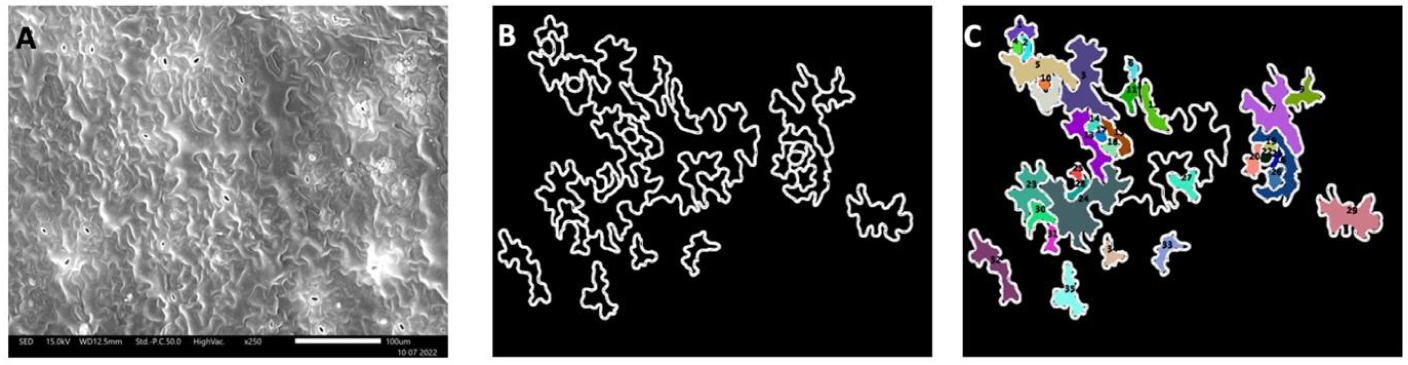
Workflow used to segment pavements cells. Using a SEM image (A), pavement cell delimitations were drawn using a graphic tablet (B). These hand-labeled images were then segmented using PaCeQuant (C) to extract cell features.

**Figure S2.**
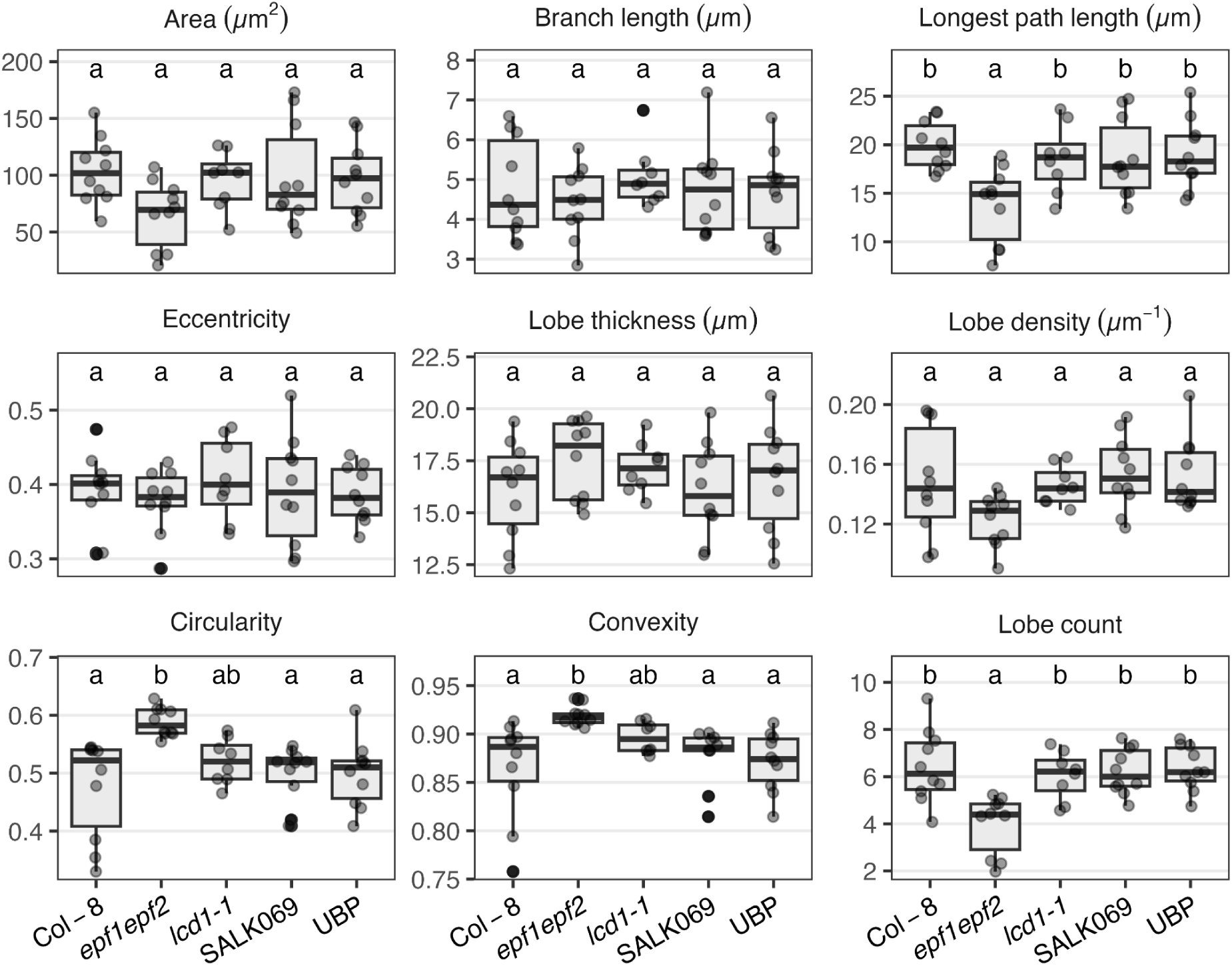
Anatomical characteristics of epidermal cells on the adaxial leaf surface in the five lines of *A. thaliana*. Different letters represent a statistically significant difference between the lines for each trait analyzed (n = 10).

**Figure S3.**
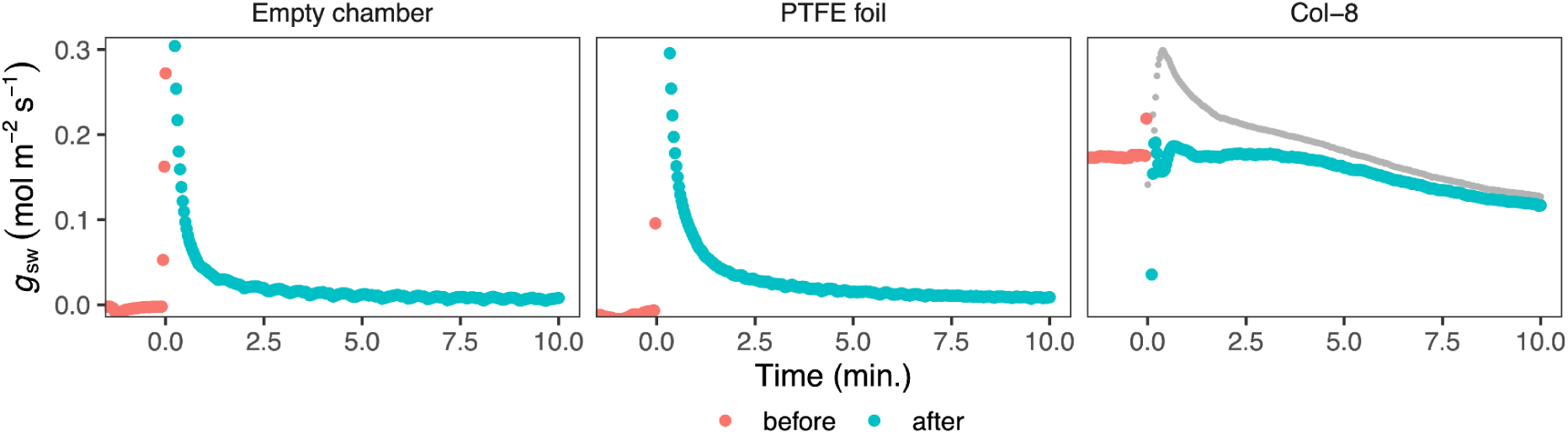
Stomatal conductance to water vapor (*g*_sw_) over time following a change in VPD from 1.2 kPa to 2.5 kPa for an empty gas-exchange chamber, for a chamber containing a 200 µm thick polytetrafluoroethylene (PTFE) foil (n=6) and for a chamber containing Col-8 leaves (n=12). Red points indicate measurements before the humidity switch. Cyan points indicate measurements after changing chamber humidity. The grey line indicates the raw measurements without any correction for mixing and water desorption (see material and methods).

**Figure S4.**
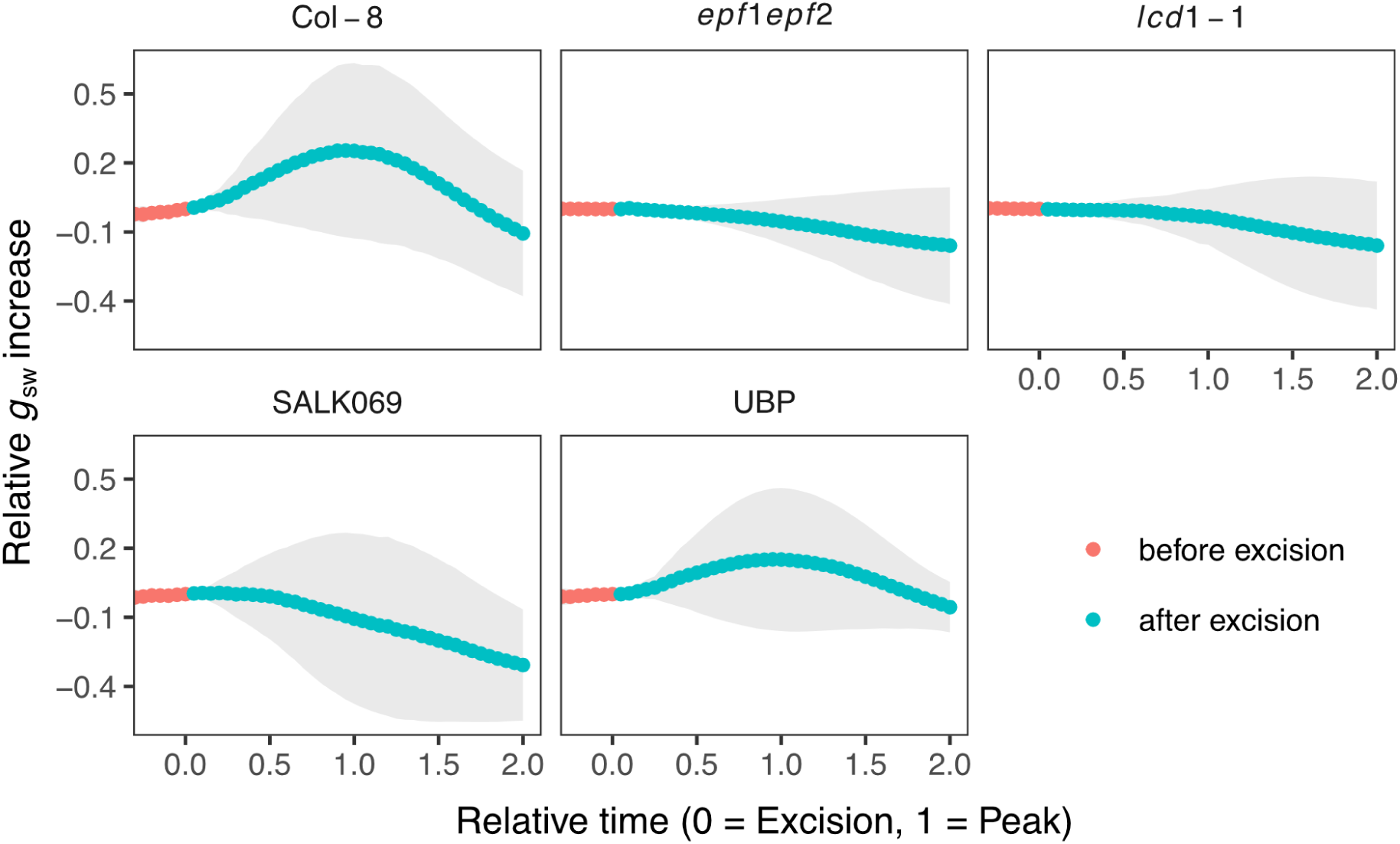
Stomatal conductance to water vapor (*g*_sw_) relative to the value at the moment of excision in five Arabidopsis lines following leaf excision. To adjust for lag time differences, relative *g*_sw_ was mapped onto a normalized response time scale, where time = 0 corresponds to the moment of excision or VPD change and time = 1 corresponds to the fitted peak of the response. For traces without a detectable peak, the Col-8 median lag time was used for normalization (262 s). Points represent mean *g*_sw_ interpolated onto a regularized 10 s grid; shaded ribbons indicate SD.

**Figure S5.**
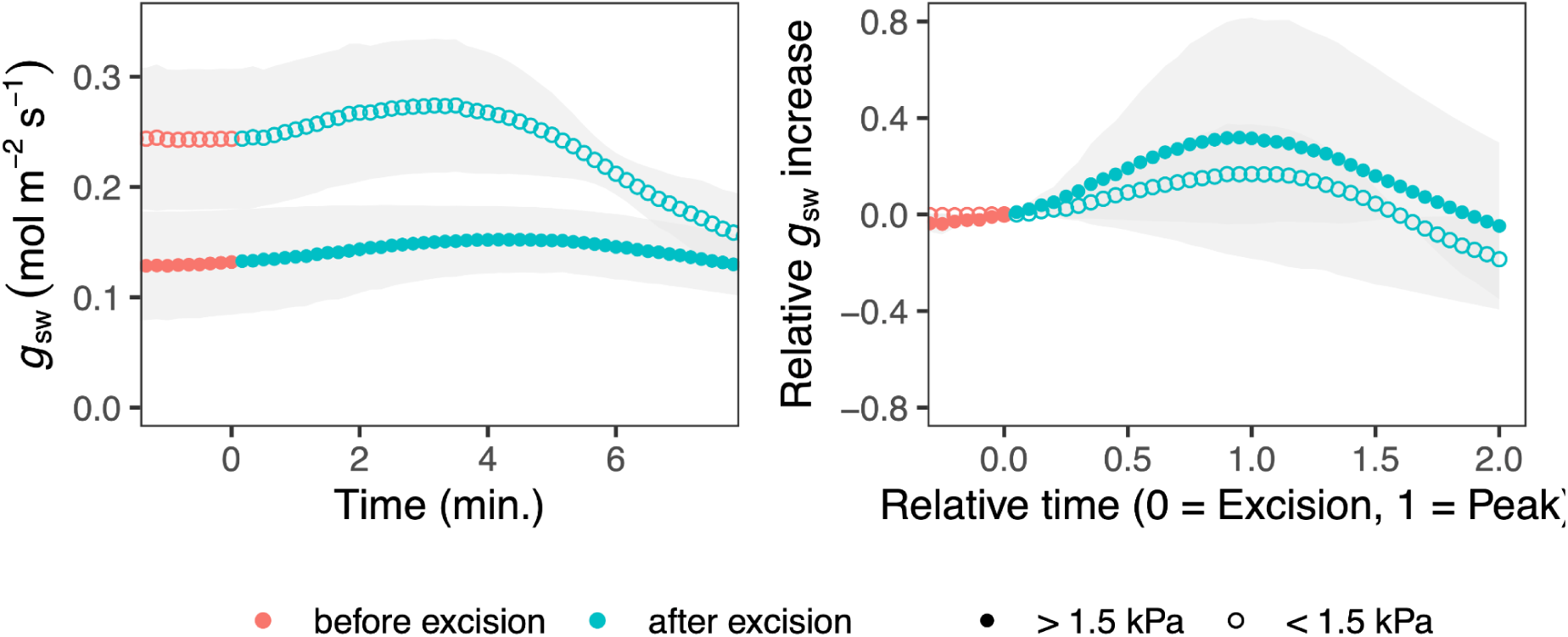
Comparison of the stomatal conductance (*g*_sw_) kinetics of excision under low VPD conditions (mean VPD before excision of 1.0 kPa, open symbols) or high VPD conditions (mean VPD before excision of 2.1 kPa, open symbols) in the Col-8 wild type *Arabidopsis thaliana*. Shown on the left as raw *g*_sw_ over time and on the right as the increase in *g*_sw_ relative to its value at the moment of excision. To adjust for lag time differences, relative *g*_sw_ was mapped onto a normalized response time scale, where time = 0 corresponds to the moment of excision or VPD change and time = 1 corresponds to the fitted peak of the response. Points represent mean values interpolated onto a regularized 10 s grid; shaded ribbons indicate SD.

**Figure S6.**
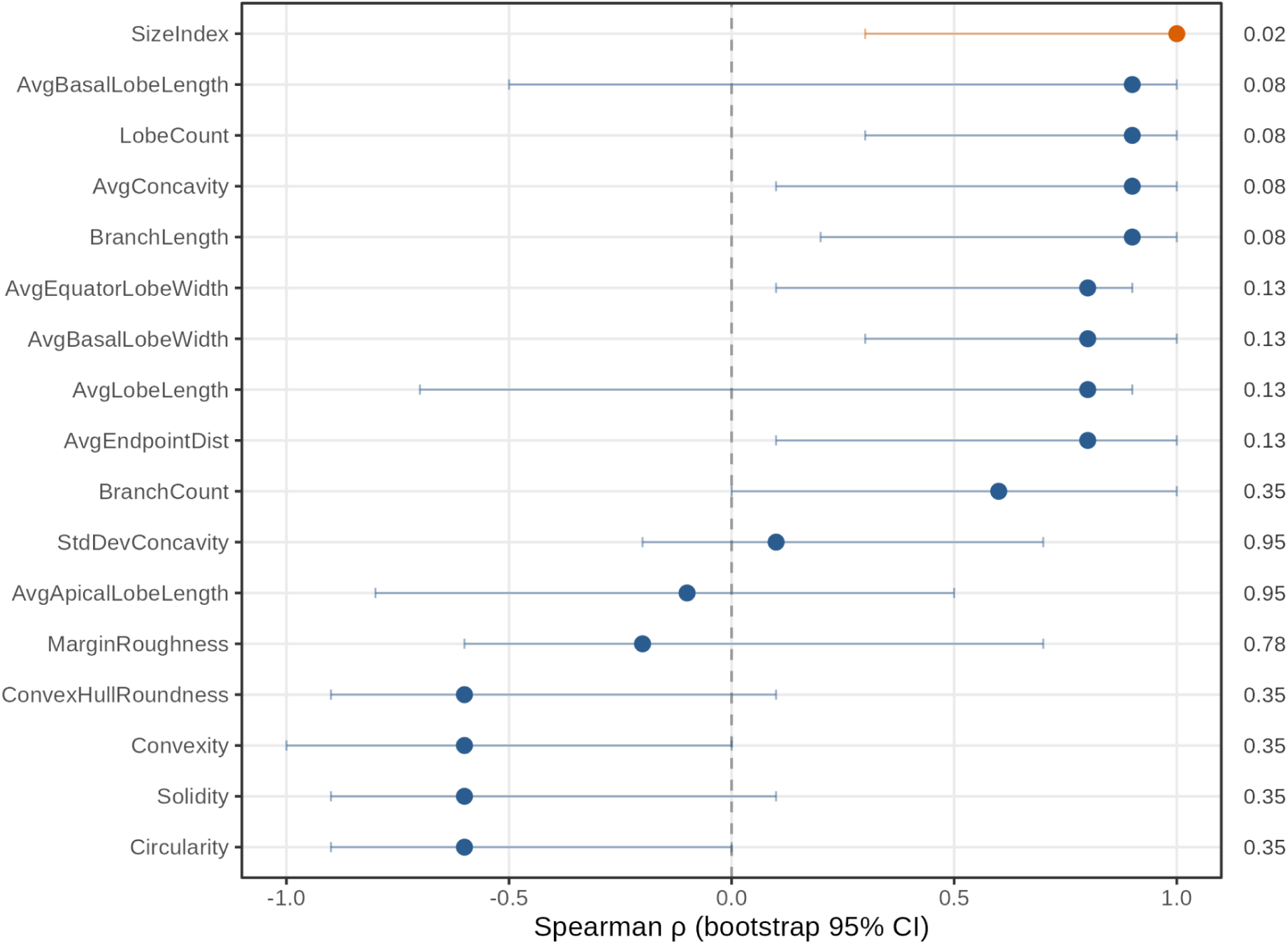
Association between pavement-cell anatomy and WWR amplitude. Points show Spearman’s ρ (inclusive bootstrap 95 % CI) for the genotype-level correlation between the relative WWR amplitude (S) and each pavement-cell trait for the abaxial epidermis. Adjacent numbers are uncorrected exact-permutation *p*-values. SizeIndex (orange) is a sign-aligned z-score composite of 11 size traits not shown individually (Area, Length, Width, LongestPathLength, ConvexHullArea, Perimeter, ConvexHullPerimeter, MinCoreWidth, MaxCoreWidth, LargestEmptyCircle, NonLobeArea). All other traits (blue) are single PaCeQuant descriptors: LobeCount, number of lobes; BranchCount, number of skeleton branches; BranchLength, total branch (lobe) length; AvgLobeLength, mean lobe length; AvgApicalLobeLength / AvgBasalLobeLength, mean length of apical / basal lobes; AvgBasalLobeWidth, mean lobe width at the base; AvgEquatorLobeWidth, mean lobe width at mid-length (shown as "Lobe thickness" in Figs 4 and S2); AvgEndpointDist, mean distance between neighbouring lobe tips; Circularity, 4π·Area/Perimeter^2^; Solidity, Area/ConvexHullArea; Convexity, ConvexHullPerimeter/Perimeter; ConvexHullRoundness, roundness of the convex hull; MarginRoughness, degree of margin waviness; AvgConcavity / StdDevConcavity, mean and variability of margin indentation depth. Additional details in: Möller, B., Poeschl, Y., Plötner, R. & Bürstenbinder, K. (2017) PaCeQuant: A tool for high-throughput quantification of pavement cell shape characteristics. Plant Physiology, 175(3): 998–1017.

**Figure S7.**
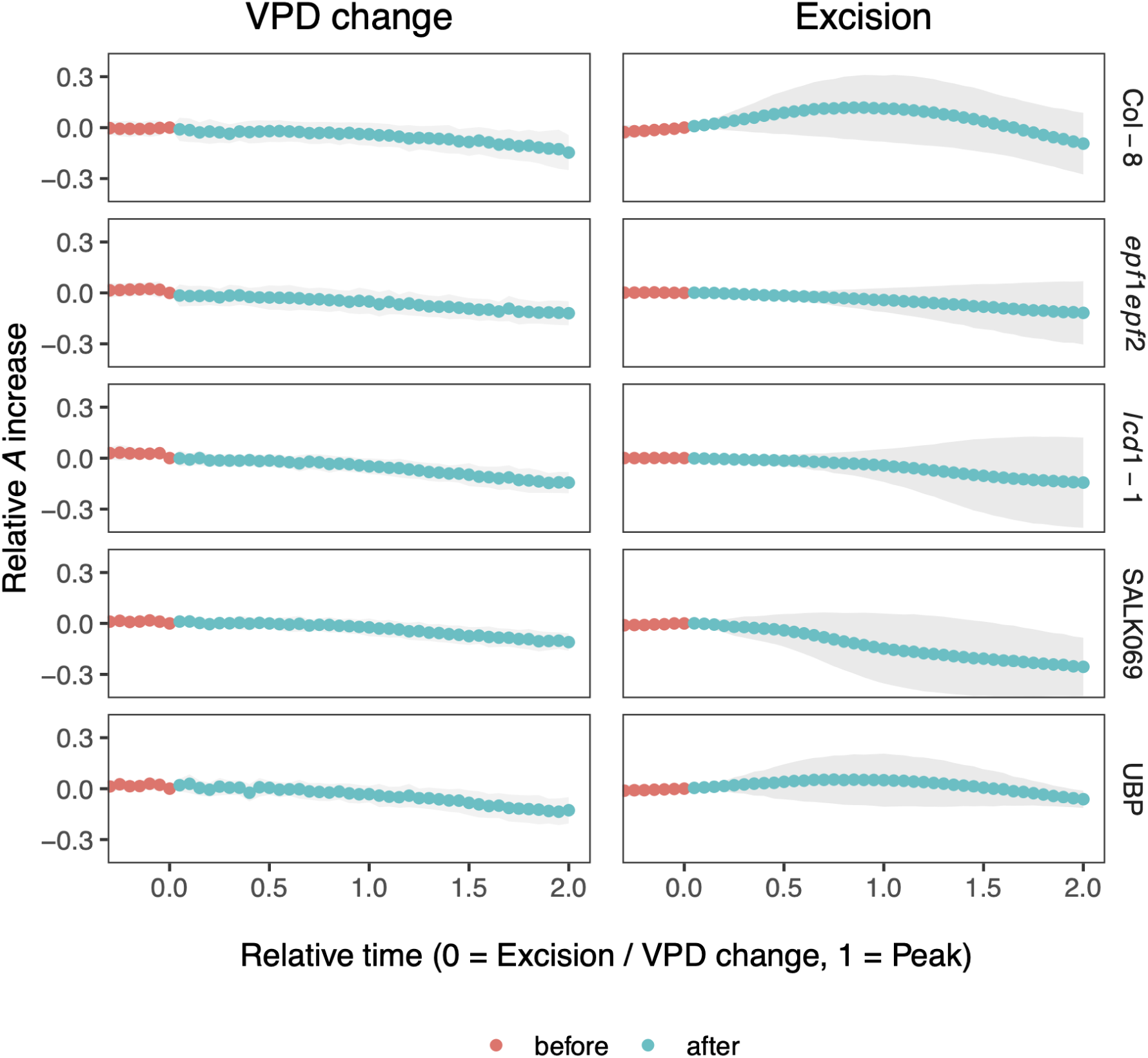
Comparison of the relative increase in photosynthetic rate (*A*) following leaf excision (left) or a step change in VPD (right). In both panels, *A* is expressed relative to its value at the moment of excision or VPD switch and mapped onto a normalized time scale, where time = 0 corresponds to the moment of excision or VPD change and time = 1 corresponds to the fitted peak of the response. For traces without a detectable peak, the excision Col-8 median lag time was used for normalization (248 s). Points represent mean values interpolated onto a regularized grid; shaded ribbons indicate ± 1 SD.

**Figure S8.**
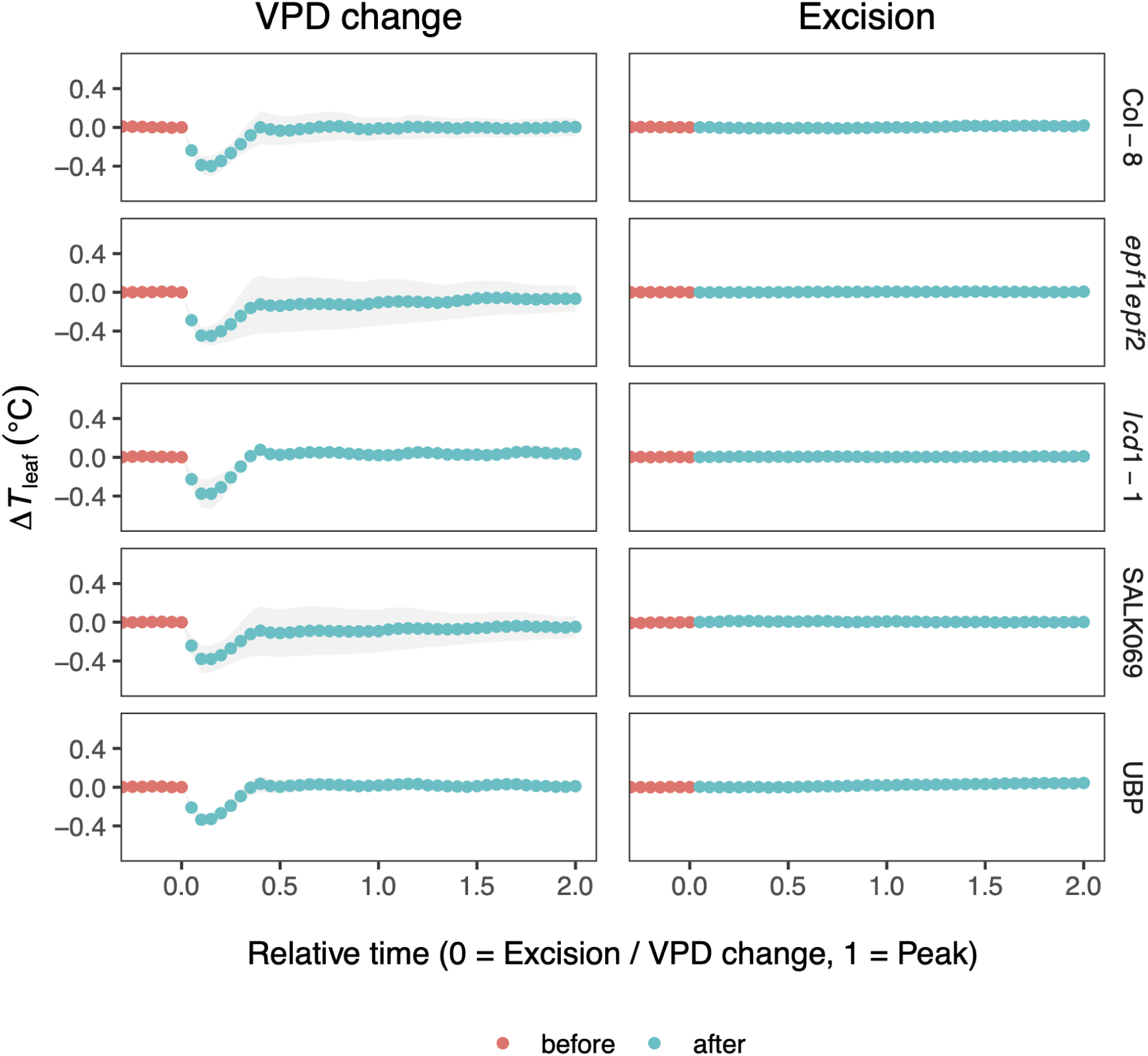
Comparison of the relative increase in stomatal conductance (*g*_sw_), photosynthetic rate (*A*), and the absolute change in leaf temperature (Δ*T*_leaf_) following leaf excision (top row) or a step change in VPD. *g*_sw_ and *A* are expressed relative to their values at the moment of excision or VPD switch, and both *g*_sw_ and *A* are expected to increase during the WWR (see also Supplemental Figure S8). For our own data, sorption-corrected values are shown for *g*_sw_ (the grey line indicates the raw measurements). Points represent mean values interpolated onto a regularized grid.

