## Supplementary figures and images for "Beyond guard cells: Epidermal anatomy determines transient stomatal opening following rapid changes in hydration status"

### Anatomical characteristics of epidermal cells on the adaxial leaf surface in the five lines of A. thaliana. Different letters represent a statisticall

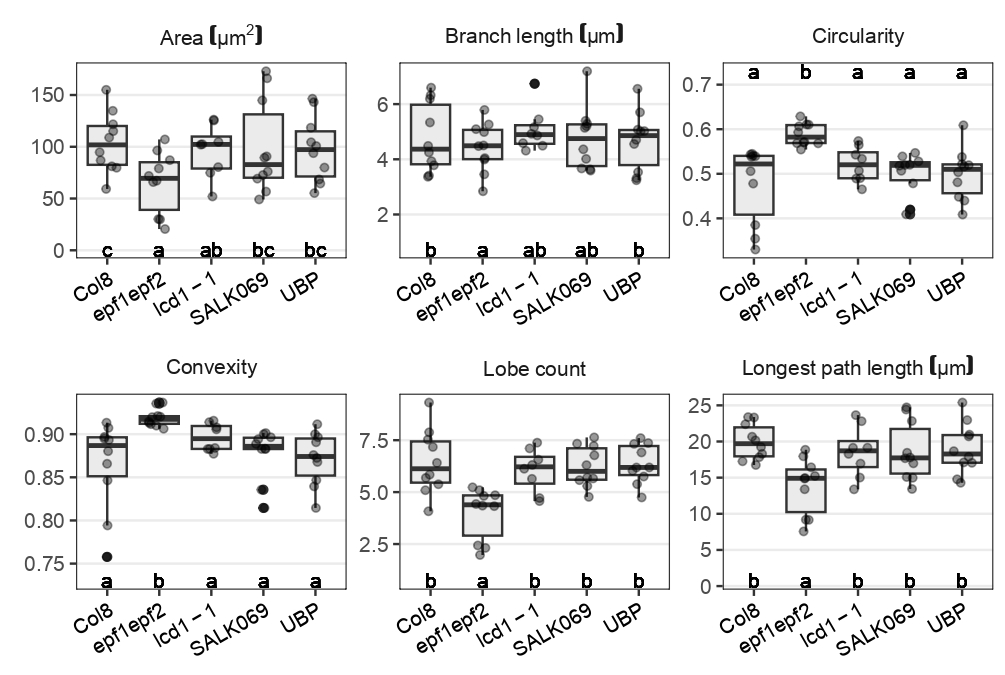

### Stomatal conductance to water vapor (gsw) over time following a change in VPD from 1.2 kPa to 2.5 kPa for an empty gas-exchange chamber, for a chamber

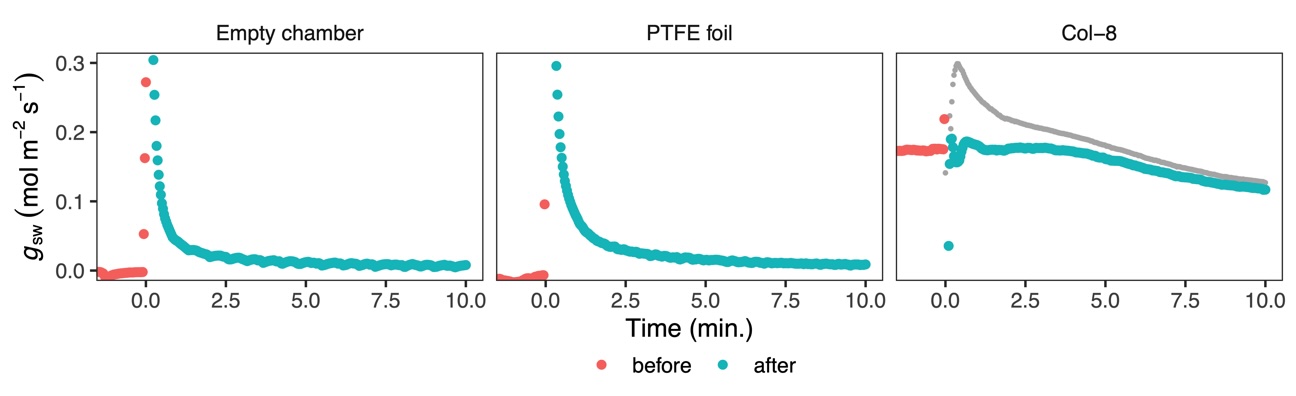

### Stomatal conductance to water vapor (gsw) relative to the value at the moment of excision in five Arabidopsis lines following leaf excision. To adjust

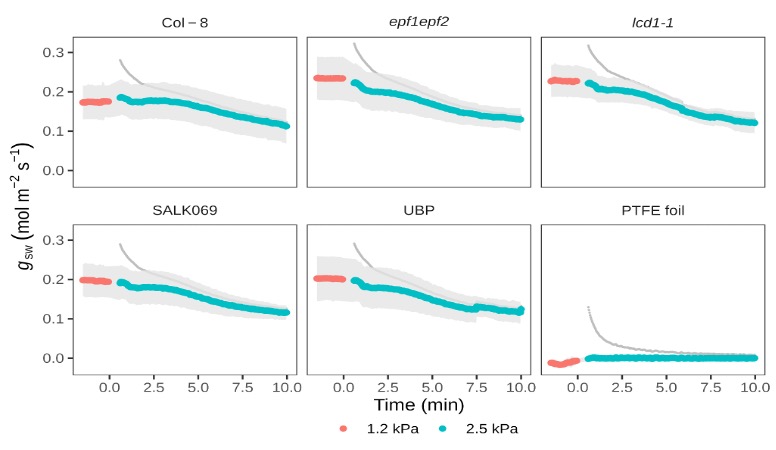

### Workflow used to segment pavements cells. Using a SEM image (A), pavement cell delimitations were drawn using a graphic tablet (B). These hand-labeled

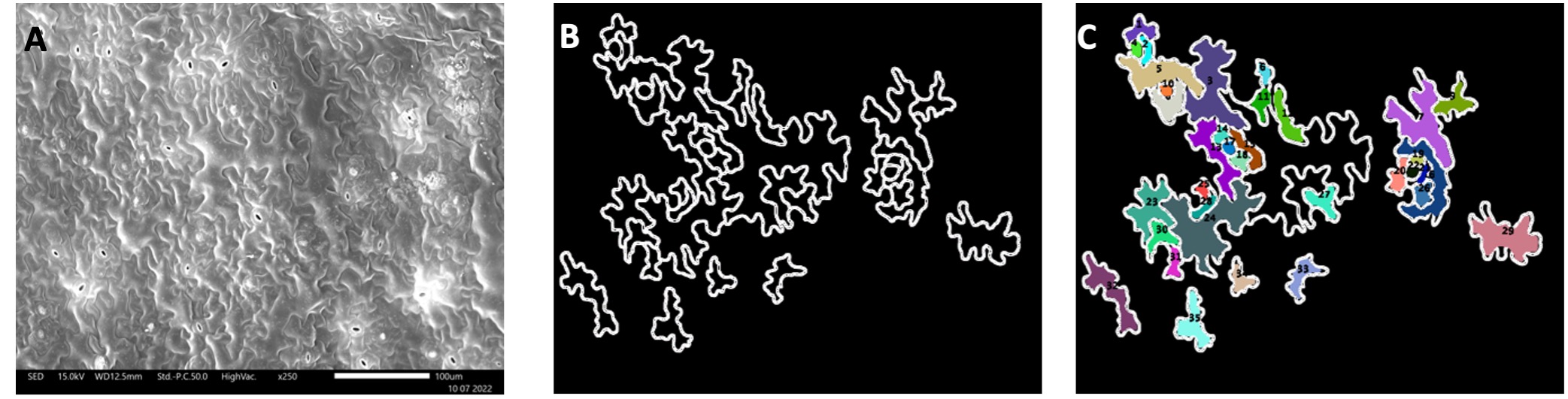
